# Patient-specific cell communication networks associate with disease progression in cancer

**DOI:** 10.1101/2021.02.08.430343

**Authors:** David L Gibbs, Boris Aguilar, Vésteinn Thorsson, Alexander V Ratushny, Ilya Shmulevich

**Affiliations:** Institute for Systems Biology, 401 Terry Avenue North, Seattle, WA 98109, USA; Bristol-Myers Squibb, 400 Dexter Avenue North, Suite 1200, Seattle, WA 98109, USA

**Keywords:** Networks, cell communication, immuno-oncology, computational oncology, bioinformatics, systems biology

## Abstract

The maintenance and function of tissues in health and disease depends on cell-cell communication. This work shows how high-level features, representing cell-cell communication, can be defined and used to associate certain signaling ‘axes’ with clinical outcomes. Using cell-sorted gene expression data, we generated a scaffold of cell-cell interactions and define a probabilistic method for creating per-patient weighted graphs based on gene expression and cell deconvolution results. With this method, we generated over 9,000 graphs for TCGA patient samples, each representing likely channels of intercellular communication in the tumor microenvironment. It was shown that particular edges were strongly associated with disease severity and progression, in terms of survival time and tumor stage. Within individual tumor types, there are predominant cell types and the collection of associated edges were found to be predictive of clinical phenotypes. Additionally, genes associated with differentially weighted edges were enriched in Gene Ontology terms associated with tissue structure and immune response. Code, data, and notebooks are provided to enable the application of this method to any expression dataset (https://github.com/IlyaLab/Pan-Cancer-Cell-Cell-Comm-Net).

## Introduction

The maintenance and function of tissues depends on cell-cell communication (Wilson et al., 2000; Haass and Herlyn, 2005). While cell communication can take place through physically binding cell membrane surface proteins, cells also release ligand molecules that diffuse and bind to receptors on other cells (paracrine or endocrine), or even the same cell (autocrine), triggering a signaling cascade that can potentially activate a gene regulatory program (Cameron and Kelvin, 2013; Heldin et al., 2016; Cohen and Nelson, 2018). More generally, a message is sent and received, transferring some information as part of a large network (Frankenstein et al., 2006). Cells communicate in order to coordinate activity, such as, to correctly (and jointly) respond to environmental changes (Song et al., 2019).

Altered cellular communication can cause disease, and conversely diseases can alter communication (Wei et al., 2004). Cancer, once thought of as purely a disease of genetics, is now recognized as being enmeshed in complex cellular interactions within the tumor microenvironment (TME) (Trosko and Ruch, 1998). The cell-cell interactions are important for cell differentiation, tumor growth (West and Newton, 2019), and response to therapeutics (Kumar et al., 2018).

Between cells, information transfer is directional in nature, where cells produce molecules that are received by the properly paired, and expressed, receptor. There is often a sender and receiver, which makes the cell-cell networks directionally linked by molecules. The dynamics of the signal is greatly important (Fridman et al., 2012, Behar et al., 2013), but unfortunately is difficult to detect in bulk sequencing experiments. One approach to studying cell interactions is through the use of graphical models of communication networks (Morel et al., 2017). By incorporating experimental data, the graphical models can become quantitative, providing predictions that can be tested and used in discovering novel drug targets and developing optimal intervention strategies.

In recent work (Thorsson et al., 2018), we developed a method used to identify cellular communication networks at work in the tumor microenvironment. Given a set of samples with a similar tumor microenvironment, the method identified ligands, receptors and cells meeting certain criteria of abundance and concordance within that set of samples. The method was applied to identify networks playing a role within specific tumor types and molecular subtypes and is available as a workflow and interactive module on the iAtlas portal for immuno-oncology (Eddy et al., 2020).

In this work, we have combined multiple sources of data with a new probabilistic method for constructing *patient-specific* cell-cell communication networks (Figure 1). In total, we built networks for 9,234 samples in The Cancer Genome Atlas (TCGA), starting from a network of 64 cell types and 1,894 ligand-receptor pairs. This is a rich feature set from which to investigate biological alterations in cell communication within the tumor microenvironment. We identified informative network features that are associated with disease progression. The method can be applied to any cancer type, but in this manuscript we focus on a selection of cancer types with very high mortality rates, including pancreatic adenocarcinoma (PAAD), melanoma (SKCM), lung (LUSC), and cancers of the gastrointestinal tract (ESCA, STAD, COAD, READ) (Cancer Genome Atlas Network, 2015).

**Figure 1.**
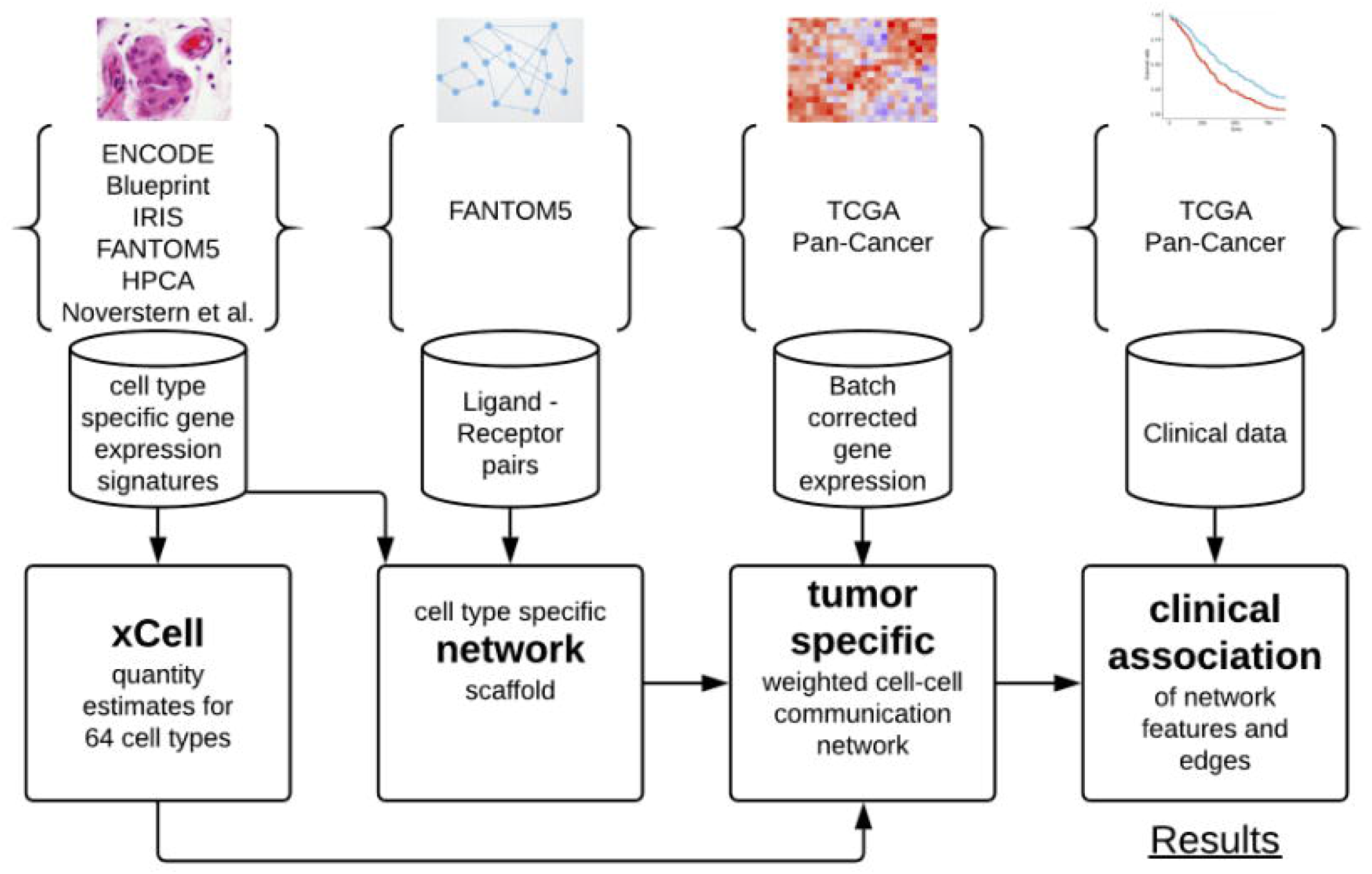
Overview of workflow showing the transition from data sources to results.

This represents a new method that provides information on possible modes of intercellular signaling in the TME, something that is currently lacking. While there are many methods on gene set scoring, cellular abundance estimation, differential expression, there are still few ways to investigate cell-cell communication diversity in the TME with respect to patient outcomes. Fortunately, new databases of receptor-ligand pairs are becoming available, making work in this area possible (Efremova et al., 2019; Jin et al., 2020; Nath and Leier, 2020; Shao et al., 2020). The methods, code, data, and complete results are available and open to all researchers (https://github.com/IlyaLab/Pan-Cancer-Cell-Cell-Comm-Net).

## Methods

### Data aggregation and integration

Data sources including TCGA and cell-sorted gene expression, bulk tumor expression, cell type scores, cell-ligand and cell-receptor presence estimations were used for network construction and probabilistic weighting on a per-sample basis.

Each tumor sample is composed of a mixture of cell types including tumor, immune, and stromal cells. Recently, methods have been developed to ‘deconvolve’ mixed samples into estimated fractions of cell type quantities. For example, xCell, which resembles gene set enrichment, has performed this estimation for 64 cell types across most TCGA samples (Aran et al., 2017). We use these xCell estimates of cellular fractions in this work.

Ramilowski et al. performed a comprehensive survey of cellular communication, generating a compendium that includes 1,894 ligand-receptor pairs, and a mapping between 144 cell types and expression of ligand or receptor molecules (Ramilowski et al., 2015) The compendium was shared via 5th edition of the FANTOM Project, FANTOM5. These ligand-receptor pairs were adopted for this study. Unfortunately, the FANTOM5 collection of cell types does not overlap well with cell types in xCell. In order to integrate the xCell and FANTOM5 data resources, it was necessary to determine the expressed ligands and receptors for each of the 64 cell types in xCell, using the source gene expression data.

The xCell project used six public cell sorted bulk gene expression data sets in order to generate gene signatures and score each TCGA sample. Across the data sets, there is some discrepancy in cell type nomenclature, making it necessary to manually curate cell type names to improve alignment across experiments (Supplementary Table 1). Typically, for a given cell type, there are several replicate expression profiles, often across the data sets.

### Building the cell-cell communication network scaffold

In the FANTOM5 ‘draft of cellular communication’, an expression threshold of 10 TPMs was used to link a cell type to a ligand or receptor. When considering the distribution of expression in the FANTOM5 project, 10 TPMs is close to the median.

To construct our scaffold, we used a majority voting scheme based on comparing expression levels to median levels. For each cell type, paired with ligands and receptors, if the expression level was greater than the median, it was counted as a vote (i.e., ligand expressed in this cell type). If a ligand or receptor recieved a majority vote across all available data sources, it was accepted, and entered into the cell-cell scaffold.

With this procedure, a network scaffold is induced, where cells produce ligands that bind to receptors on receiving cells. One edge in the network is composed of components cell - ligand - receptor - cell. This produced a cell-cell communication network with over 1M edges. Each edge represents a possible interaction in the tumor microenvironment. We subsequently determine the probability that an edge is active in a particular patient sample using a probabilistic method described below.

### Patient level cell-cell communication network weights

With a cell-cell scaffold, expression values and cell type estimations per sample, we can produce a per-sample weighted cell-cell communication network (Figure 2). This is done probabilistically, using the following definition:

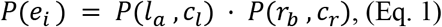

where *e_i_* is edge *i*, *l_a_* is ligand *a*, *r_b_* is receptor *b,* and *c_l_* and *c_r_* are cells that can produce ligand *a* and receptor *b* respectively. *P*(*e_i_*) represents a probability that edge *i* is active and is based on the premise that the physical and biochemical link and activation is possible only if all the components are present, and that activity becomes increasingly possible with greater availability of those components. The joint probabilities can be decomposed to:

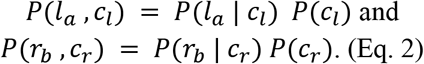

**Figure 2.**
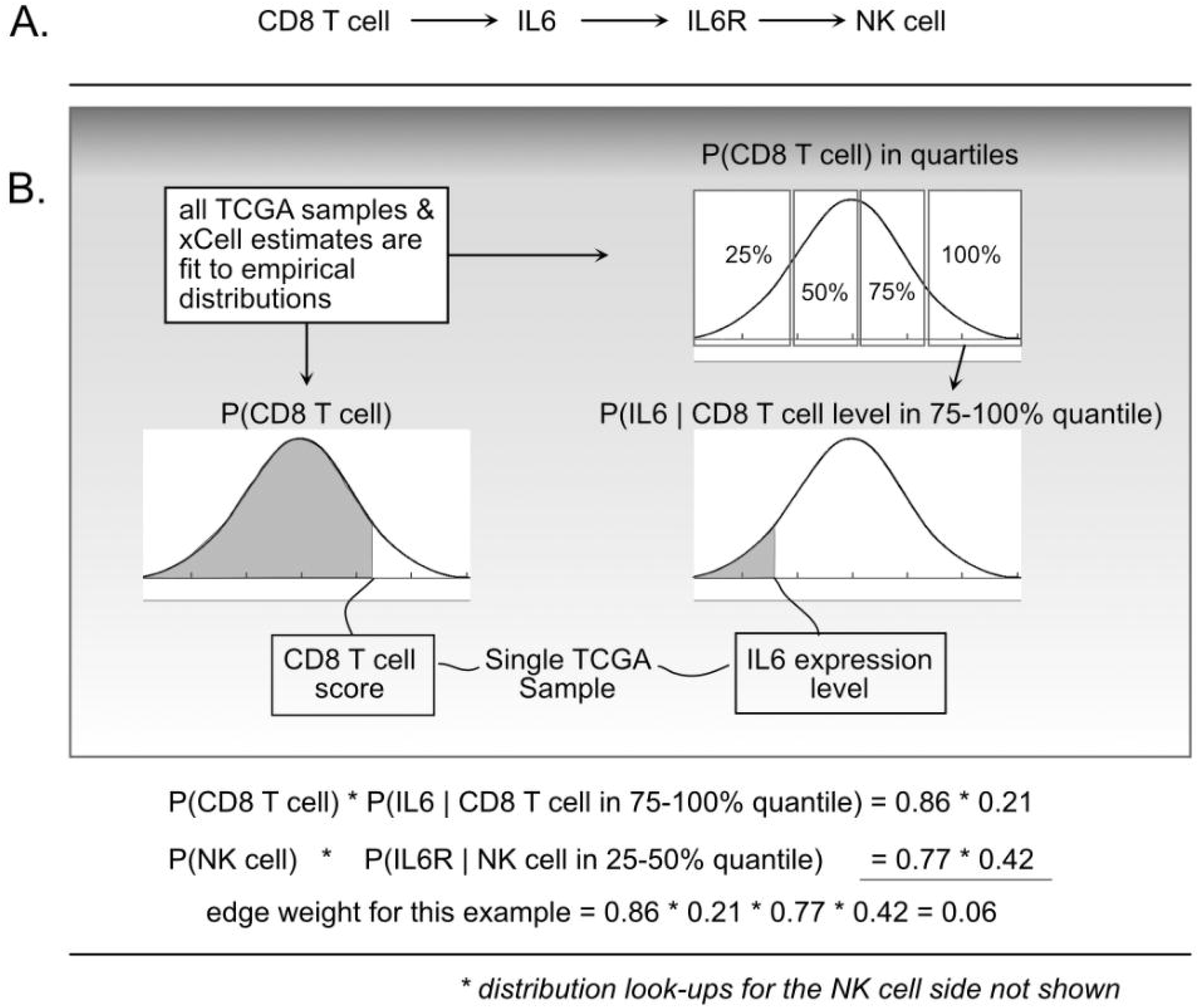
Illustration of the probabilistic model and edge weight computations. (A) For a given cell-cell communication edge, (B) per patient values are used to ‘look up’ probabilities from the distributions learned from all TCGA data. Those probabilities are then used to compute an edge weight.

The *P*(*c_lk_*) is short for CDF *P*(*C_l_* < *c_lk_*) which indicates the probability that a randomly sampled value from the empirical *C_l_* distribution (over all 9K TCGA samples) would be less than the cell estimate for cell type *l*, in sample *k*. To do this, for a given cell type, using all samples available, an empirical distribution *P*(*C_l_*) is computed, and for any query, essentially using a value *c_lk_*, the probability can be found by integrating from 0 to *c_lk_*.

To compute *P*(*l_a_* | *c_l_*), each *C_l_* distribution was divided into quartiles, and then (again using the 9K samples) empirical gene expression distributions within each quartile were fit. This expresses the probability that with an observed cell quantity (thus within a quartile), the probability that a randomly selected gene expression value (for gene *l_a_*) would be lower than what is observed in sample *k.*

We refer to “edge weights” to be the probability as shown in Eq. (1). To compute edge weights, each TCGA sample was represented as a column vector of gene expression and a column vector of cell quantities (or enrichments). For each edge in the scaffold (cell-ligand-receptor-cell), data was used to look up probabilities using the defined empirical distributions and taking products for the resulting edge weight probability. This leads to over 9K tumor-specific weighted networks, one for each TCGA participant.

Probability distributions were precomputed using the R language empirical cumulative distribution function (ecdf). For example, fitting *P*(CD8 T cells) is done by taking all available estimates across the Pan-Cancer samples and computing the ecdf. Then, for a sample *k,* we find *P*(*C_l_* < *c_lk_*) using the ecdf. The same technique is used to find the conditional probability functions, where for each gene, the expression values are selected after binning samples using the R function ‘quantile’, and then used to compute the ecdf. With all distributions precomputed, 9.8 billion joint probability functions were computed using an HPC environment, then transferred to a Google BigQuery table where analysis proceeded. This table of network weights was structured so that each row contained one weight from one edge and one tumor sample. Although being a large table of 9.8 billion rows, taking nearly 500GB, BigQuery allows for fast analytical queries that can produce statistics using a selection of standard mathematical functions.

### Association of network features and survival-based phenotypes

The *S_1_* statistic is a robust measure based on the difference of medians (Yahaya et al., 2004; Ahad et al., 2016; Babu et al., 1999; Hubert et al. 2012), in this case the median of edge weights for a defined phenotypic group. *S_1_* statistics were computed using the NCI cancer research data commons cloud resource, the ISB-CGC, per tissue type.

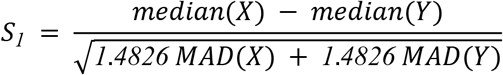

This statistic allowed for cell-cell interactions to be ranked within a defined context. The results were again saved to BigQuery tables to allow for further cloud-based analysis and integration with underlying data.

To judge the magnitude of the statistic with respect to a random context (Figure 3), an ensemble of three edge-weight sample-pools were generated, each with 100K weights. Then, for each member of the ensemble, 1 million *S_1_* statistics were generated using sample sizes that match the analyzed data. These random *S_1_* statistic distributions were used to compare to the observed results (i.e., a resampling procedure).

**Figure 3.**
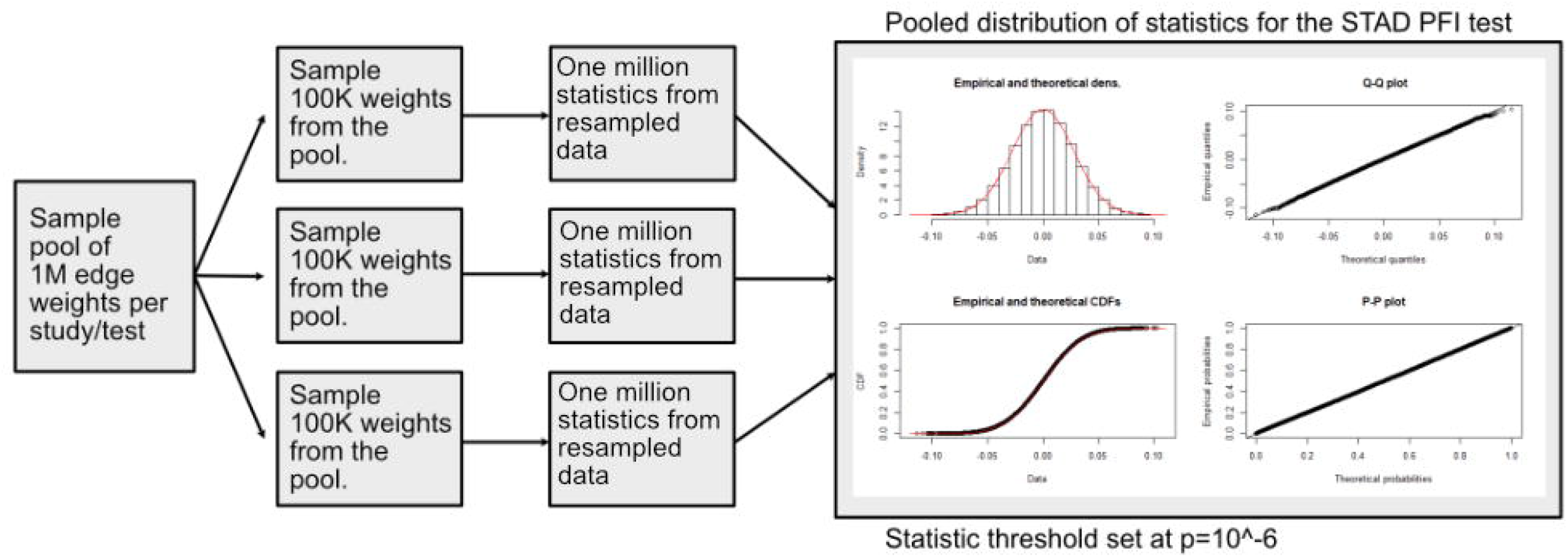
Diagram of how differentially weighted edges were determined. Three samples of edge weights were taken from the pool by tissue source. Then matching the sample proportions in the clinical features, permutations were sampled and used for computing randomized *S_1_* statistics. Each sample was used to produce 1 million permuted statistics, and taken together, the millionth percentile was used as a cutoff in determining important edges.

As an initial examination of the interplay of cell communication and disease, two proxies of disease severity were investigated: progression-free interval (PFI) and tumor stage (Liu et al., 2018). The staging variable used the AJCC pathologic tumor stage. The PFI feature was computed using days until a progression event. The staging variable was binarized by binning stages I-II together (“early stage”), and III-IV together (“late stage”). A binary PFI variable was created by computing the median PFI on non-censored samples and then applying the split to all samples. Both clinical features were computed by tissue type (TCGA Study). As Liu et al writes, “The event time is the shortest period from the date of initial diagnosis to the date of an event. The censored time is from the date of initial diagnosis to the date of last contact or the date of death without disease.”

For example, in LUSC, the median time to PFI event was 420 days (14.0 months) and in the censored group, 649 days (21.1 months). After splitting samples at 420 days (14 months), the short PFI group was composed of 67 uncensored samples and 128 censored samples. The long PFI group was composed of 68 uncensored samples and 234 censored samples.

Null distributions, using these same sample sizes (e.g., one group of 68 and another group of 234), were generated by repeatedly drawing from the previously described ensemble of three sample-pools. The distributions, while heavy tailed, were close to Normal (Supplemental Figure 1). The *S_1_* statistics scale with the difference in median values (Supplemental Figure 4). After combining resampled statistics across the ensemble, an edge was selected as a high edge weight if it were in the top 1 millionth percentile when compared to the null. Each tissue and contrast generates a weighted subgraph of the starting scaffold, which is retained for further analysis (e.g., a LUSC-PFI network).

To identify informative cell-cell edges that relate to disease progression, machine learning models were trained on binarized clinical data as described. With clinical features such as progression free interval (PFI) and tumor stage for each sample, a matrix of patient-specific edge weights was constructed representing each tissue and contrast. Classification of samples was performed with XGBoost classifiers (Chen and Guestrin, 2016), which are composed of an ensemble of tree classifiers. To avoid overfitting the models, the tree depth was set at maximum of 2 and the early-stopping parameter was set at 2 rounds (training was stopped after the classification error did not improve on a test set for two rounds). XGBoost provides methods for determining the information gain of each feature in the model and was used to rank edges that are most informative for classification.

Gene ontology (GO) term enrichment was performed using the GONet tool (Pomaznoy et al., 2018). The set of 1,175 genes in the cell-cell scaffold was used as the enrichment background. GONet builds on the “goenrich” software package, which maps genes onto terms and propagates them up the GO graph, performs Fisher’s exact tests, and moderates results with FDR. To compare the results, random collections of genes were generated from the cell-cell scaffold and produced no significant results.

## Results

The scaffold network graph is heterogeneous, containing nodes representing cells, ligands (e.g. cytokines), and receptors. Edges are directed, following communication routes from cell to cell. But, to simplify the graph, a cell produces a ligand that binds a receptor found on another cell type, which could make a single edge “LCell-Ligand-Receptor-RCell”. In total, there were 1,062,718 cell-cell edges in the network.

The number of edges for ligand-producing cells varies from 32,910 for osteoblasts to 6,587 for Multi-Potent Progenitor (MPPs). For receptor-producing cells, the range spans from 30,225 for platelets to 5,763 edges for MPPs.

Applying the proposed probabilistic framework allowed for the creation of 9,234 weighted networks. The edge weight distributions generally follow approximately exponentially decreasing function (Supplemental Figure 1). There are few edges with strong weights and many with low (near zero) weights.

We first sought to find communication edges that were most characteristic of an individual tumor type. The *S_1_* statistics comparing one tumor type to all other tissue types was computed, with a high score indicating a substantial difference in edge weights between the two groups. Edges were found that clearly delineated tissues (Figure 4). For example, in SKCM (skin cutaneous melanoma), the top scoring edge is between melanocytes the most cell of origin for cutaneous melanoma (Melanocytes-MIA-CDH19-Melanocytes, *S_1_* score 2.5, median edge weight 0.86 higher than in other tumor types). Normal tissue differences can contribute to differences in edge weights, though in this case the central role of melanocytes in melanomas implies that the high scores are likely due to cancerous cell activity. The study with the most similar edge weights is uveal melanoma (UVA), which arises from melanocytes resident in the uveal tract (Robertson et al., 2017) (Fig. 4A). Additionally, we observed that when a cell type is highly prevalent in a particular tissue, and the scaffold has an autocrine loop, interactions between that type of cell tend to have high weights. If we exclude cell types communicating with self-types, then for SKCM, osteoblasts, natural killer T cells, and mesenchymal stem cells (MSCs) interact with melanocytes in the top 10 scoring edges, consistent with the emerging role of these cell types in melanoma. An important role for osteoblasts is now coming to light for melanoma (Ferguson et al., 2020). Natural killer T cells are being investigated for their applicability in immunotherapy of cancers such as melanoma (Wolf et al., 2018). MSCs appear to interact with melanoma cells, as work by Zhang et al. (Zhang et al., 2017) showed the proliferation of A375 cells (a melanoma cell line) was inhibited and the cell cycle of A375 was arrested by MSCs, and cell-cell signaling related to NF-κB was down-regulated. Overall, the number of high weight edges in each tumor type did not associate with the number of samples, as might be expected (Supplemental Figure 2).

**Figure 4.**
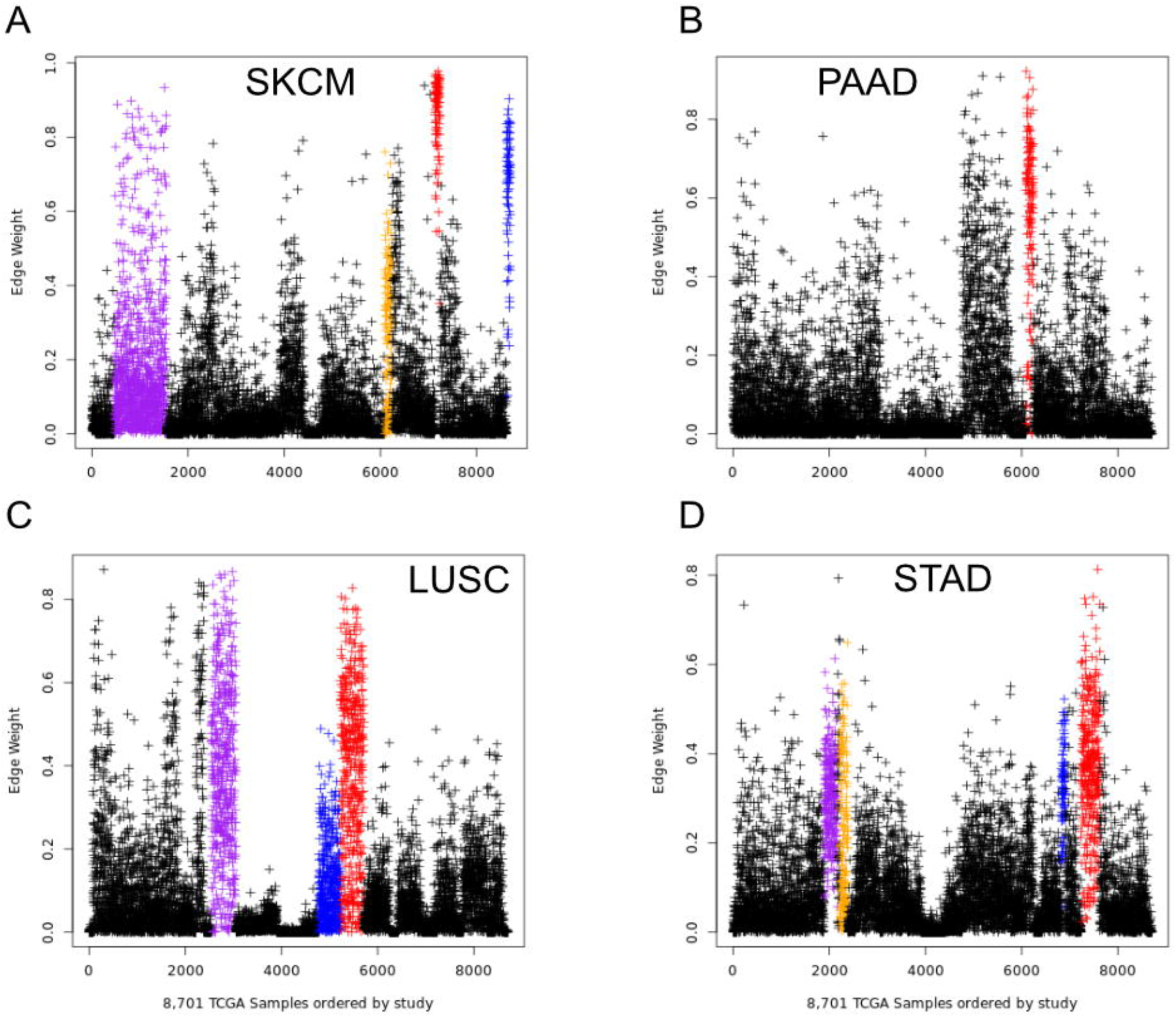
Top edges (by *S_1_* scores) that can distinguish tissue types. Each point represents a tumor sample and each panel represents one edge. (A) EdgeID 605551, Melanocytes-MIA-CDH19-Melanocyte SKCM red, UVM blue, BRCA purple, PAAD orange. (B) EdgeID 687457, MSC-TFPI-F3-MSC, PAAD red. (C) EdgeID 968128, Sebocytes-WNT5A-FZD6-Sebocytes, LUSC red, LUAD blue, HNSC purple. (D) EdgeID 1049823, Th2 cells-IL4-IL2RG-Megakaryocytes, STAD red, READ blue, COAD purple, ESCA orange.

To identify which elements of cellular communication networks might be associated with clinical progression of particular tumor types, we identified edges associated with disease. Disease progression and severity were examined using dichotomous values of tumor stage and progression free interval (PFI) as described in the methods. Statistical scores were calculated comparing edge weight distributions between the two clinical groups using *S_1_*. Results were carried forward if larger than the threshold set by the millionth percentage of resampled statistics (Supplemental Figure 3-5), yielding differentially weighted edges (DWEs).

Most tumor types showed DWEs for PFI, and fewer for the early to late tumor stage comparison (Supplemental Figure 5). For example, STAD (gastric adenocarcinoma) had several hundred edges in for both comparisons, while PAAD (pancreatic adenocarcinoma) showed fewer DWEs, and only for PFI. Figure 5 shows median edge weights between the two groups for the selected studies. Some tumor types, like SKCM, show much stronger deviations between the medians, compared to the other studies like STAD, ESCA, and LUSC, which may be an indication of a stronger immune response. According to CRI-iAtlas (Eddy et al., 2020), among our example studies, SKCM has the highest estimated level of CD8 T cells and generally has a robust immune response.

**Figure 5.**
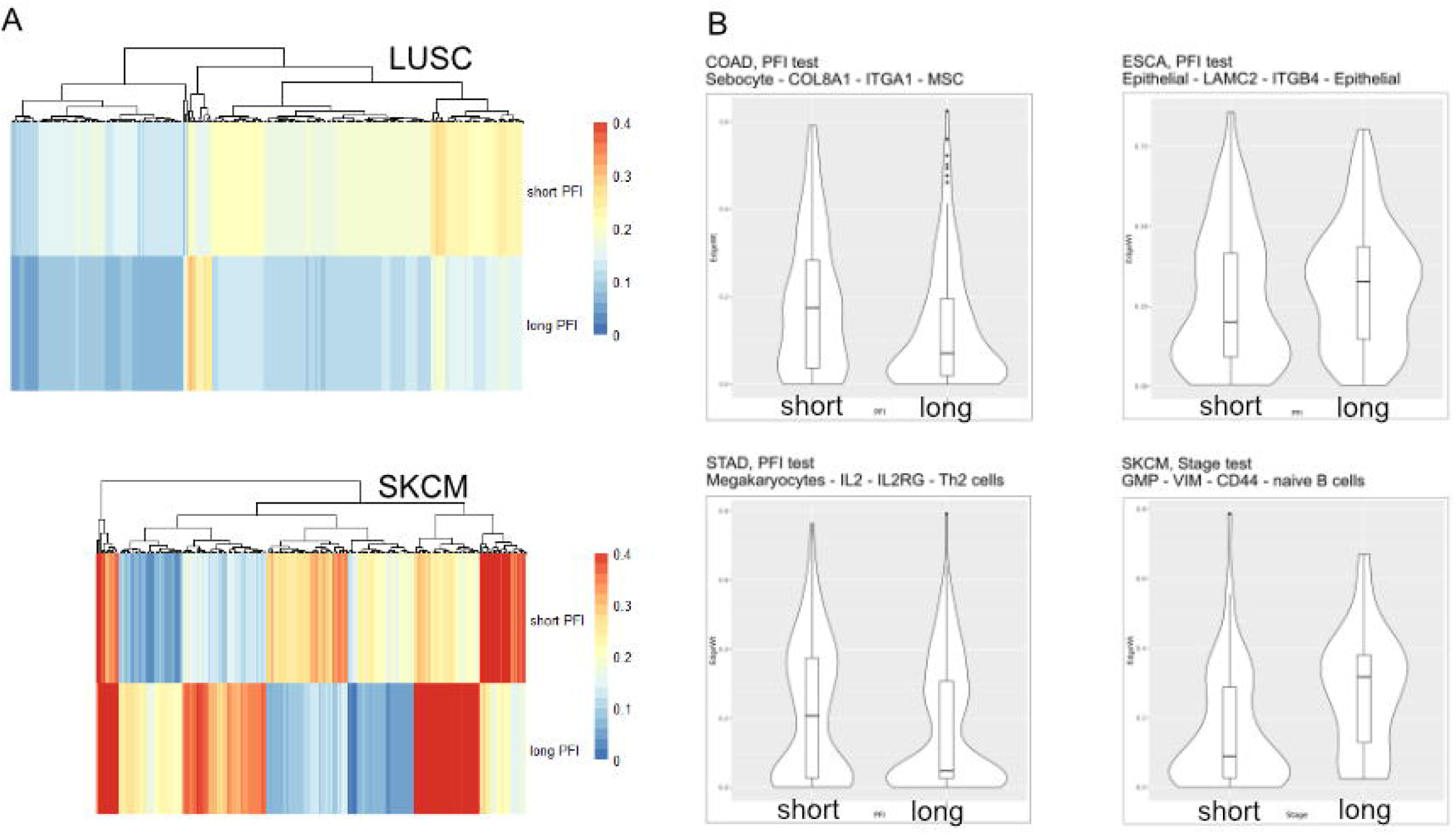
(A) Median values for each differentially weighted cell-cell edge (DWE) for the PFI categories (in row, DWE edges in columns). (B) Examples of differentially weighted edges.

Tumor stage comparison showed DWEs in 17 of 32 studies and ranged widely from 4 edges for MESO (mesothelioma) to over 63K edges for BLCA (urothelial bladder cancer adenocarcinoma). The PFI comparison showed results in 28 / 32 studies and ranged from 4 edges in READ to over 21K in LIHC. See Table 1 for edge counts from selected studies. The studies with larger numbers of samples had permuted *S_1_* distributions that were narrow compared to studies with few samples (Supp. Fig. 3), but there was not a strong association between DWE counts and sample sizes. The variation thus more likely has to do with clinical factors.

**Table 1.** Counts of differentially weighted edges compared to the number of samples in each study.

| Study | N samples | PFI short/long | PFI DWEs | Selected Feat. | Model Accuracy | GO results? |
| --- | --- | --- | --- | --- | --- | --- |
| ESCA | 170 | 73/97 | 137 | 36 | 94.7 | y |
| STAD | 409 | 155/231 | 142 | 78 | 95.1 | y |
| PAAD | 178 | 68/83 | 8 | - | - | y |
| COAD | 281 | 96/183 | 63 | 50 | 97.1 | y |
| READ | 91 | 16/71 | 4 | - | - | y |
| SKCM | 102 | 27/75 | 249 | 12 | 91.1 | y |
| LUSC | 494 | 193/285 | 304 | 119 | 98.7 | y |
| Study | N samples | Stage early/late | Stage DWEs | Selected Feat. | Model Accuracy | GO results? |
| ESCA | 170 | 86/63 | 0 | - | - | - |
| STAD | 409 | 167/198 | 241 | 114 | 99.7 | y |
| PAAD | 178 | 142/7 | 0 | - | - | - |
| COAD | 281 | 151/118 | 1851 | 84 | 99.6 | y |
| READ | 91 | 36/44 | 34 | 18 | 97.5 | y |
| SKCM | 102 | 68/29 | 221 | 8 | 99 | n |
| LUSC | 494 | 390/89 | 0 | - | - | - |

Study: tissue type, N samples: number of samples used, PFI short/long: number of samples in each group, PFI DWEs: number of differentially weighted edges, Model Accuracy: accuracy of predicting group, GO results?: if yes, significant GO enrichments.

Within a tumor type and clinical response variable, the set of high scoring edges were usually dominated by a small number of cell-types, ligands, or receptors (Figure 6, Supplemental Figure 6A,6B). For SKCM, in the tumor stage contrast, a majority of ligand-producing cells include GMP cells, Osteoblasts, MSC cells, and Melanocytes, in order of prevalence. The number of edges starting with these four cells accounts for 53% of DWEs. Certainly, melanocytes are well known in melanoma, and mesenchymal stem cells are drawn to inflammation, but the role of osteoblasts is less well documented, but still have been associated with melanoma progression (Ferguson et al., 2020).

**Figure 6.**
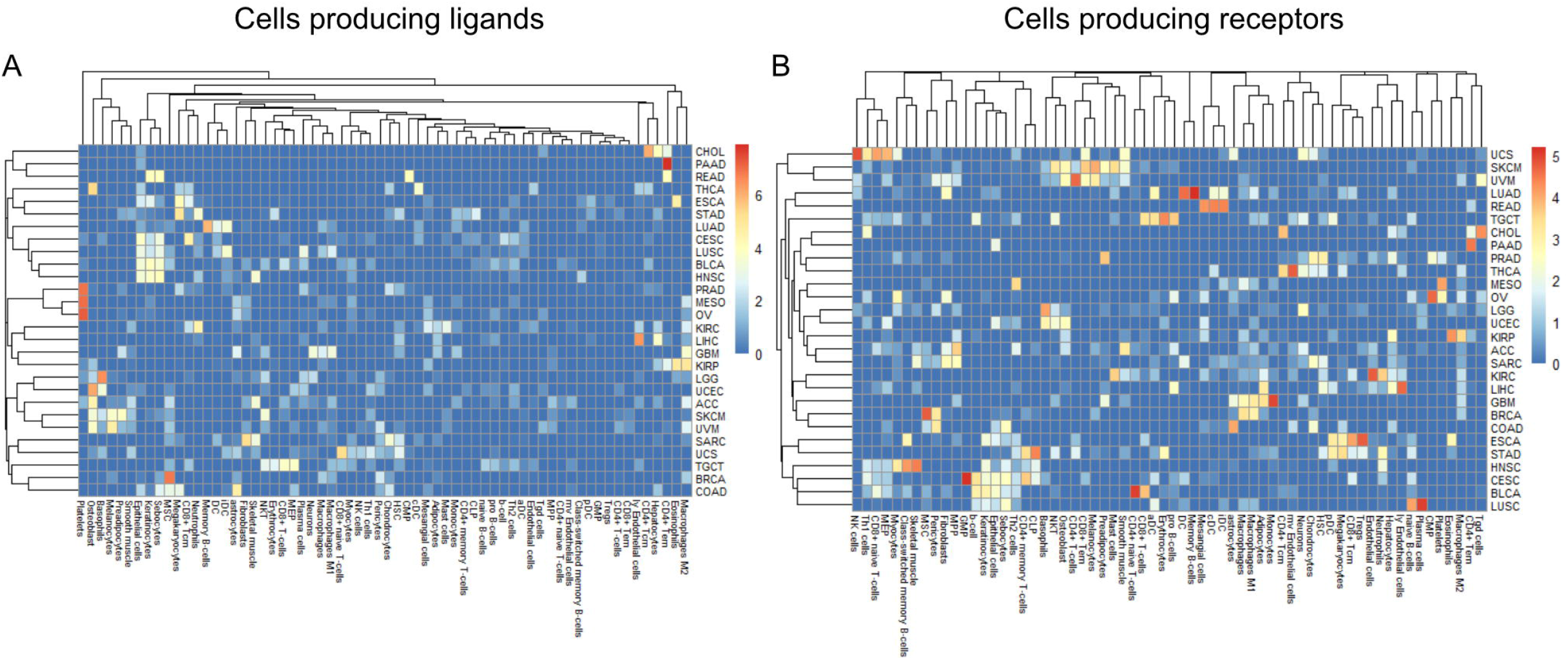
Edge member dominance in DWEs shown by log10 counts of cell types.

**Figure 7.**
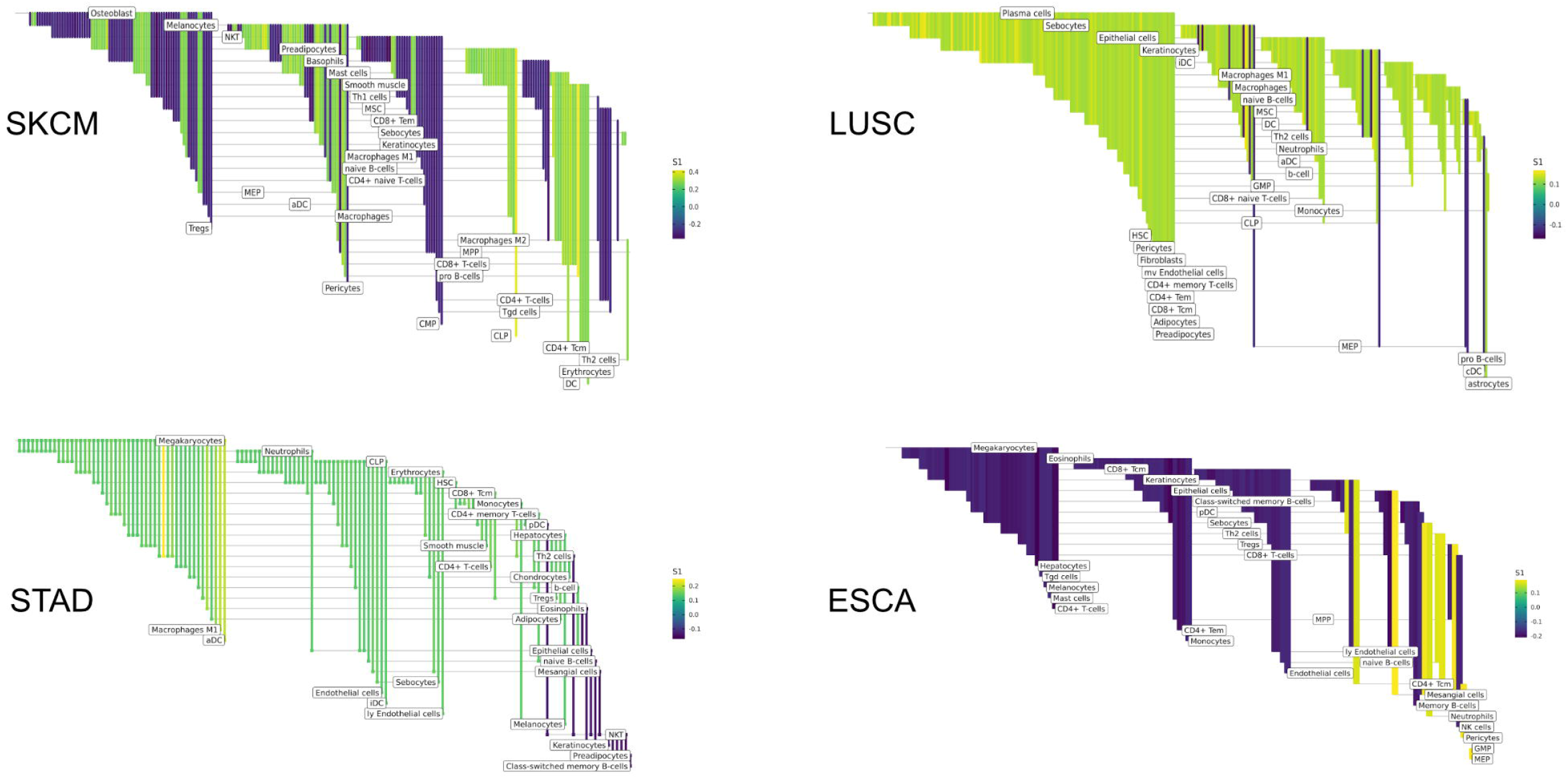
High probability edges (DWEs) from PFI contrasts form predictive connected subnetworks. Color indicates the magnitude and direction of *S_1_* statistics (+ / -).

In the PFI contrast with gastroesophageal cancers, megakaryocytes are the most common cell type in STAD DWEs (40 edges out of 142), and the second most common in ESCA (49 edges out of 137, following CD8+ Tcm interactions). The megakaryocyte DWEs include ligands and receptors that represent both interleukins and ECM-associated molecules such as integrins and collagen, but also NOTCH1 and PF4 (platelet factor 4). For STAD, most edge weights are lower with longer PFI. Put another way, the shorter PFI intervals (adverse outcome) were associated with increased megakaryocyte-involved edge weights (Supplemental Figure 7).

However, the opposite is observed in ESCA, where higher edge weights were generally associated with longer PFI (negative *S_1_* score). In ESCA, edges that show high weights for short PFI include Neutrophils-HMGB1-SDC1-Sebocytes (0.17). Although ESCA has a much lower xCell mean megakaryocyte score than STAD (38% lower), the cell score trends from xCell follow opposite trends with STAD decreasing with longer PFI and ESCA increasing with PFI. STAD is among the tissues with highest megakaryocyte scores (59, 56th rank out of 64 for PFI 1,2 resp.), ESCA is at a respectable rank of 49 and 44 out of 64 for short-PFI, respectively.

In COAD (colorectal adenocarcinoma), for ligand-producing cells, the DWEs were dominated by astrocytes, MSCs, megakaryocytes, and sebocytes, while receptor-producing cells included astrocytes, chondrocytes, and MSCs in order of counts of DWEs. By summarizing DWEs we can possibly categorize cancer types based on which cells are taking part in potentially active interactions.

The above-described edge dominance is related to cells (graph nodes) with high degree. In the language of graphs, the degree is the count of edges connected to a given node or vertex. In STAD the cell types with highest degree are megakaryocytes (degree 50), followed by neutrophils (31), CLP cells (26), and erythrocytes (23)(Supplemental Figure 6A,B). However, if we look at the directionality for the directed graph, we see that while megakaryocytes are split nearly evenly in and out, cells like the Th1 have 5 edges in, and only a single edge out, whereas B cells have zero edges in and 4 edges out. The network directionality should be considered in activities such as the modeling of dynamical systems.

Within the tumor microenvironment, communication between the multitude of cells happens simultaneously through many ligand-receptor axes. By considering a set of differentially weighted edges within a tissue type, we can construct connected networks that potentially represent dynamic communication. DWEs derived by comparing edge weights between clinical groups may indicate which parts of the cell-cell communication network shift together with disease severity.

We sought to identify which aspects of intercellular communication could relate to tumor staging or disease severity. The edges making up the differential networks were used to model clinical states for individual tumors. XGBoost models (Chen and Guestrin, 2016) were fit on each clinical feature, using edge weights as predictive variables, to infer which edges carried the most information in classification (Figure 8).

**Figure 8.**
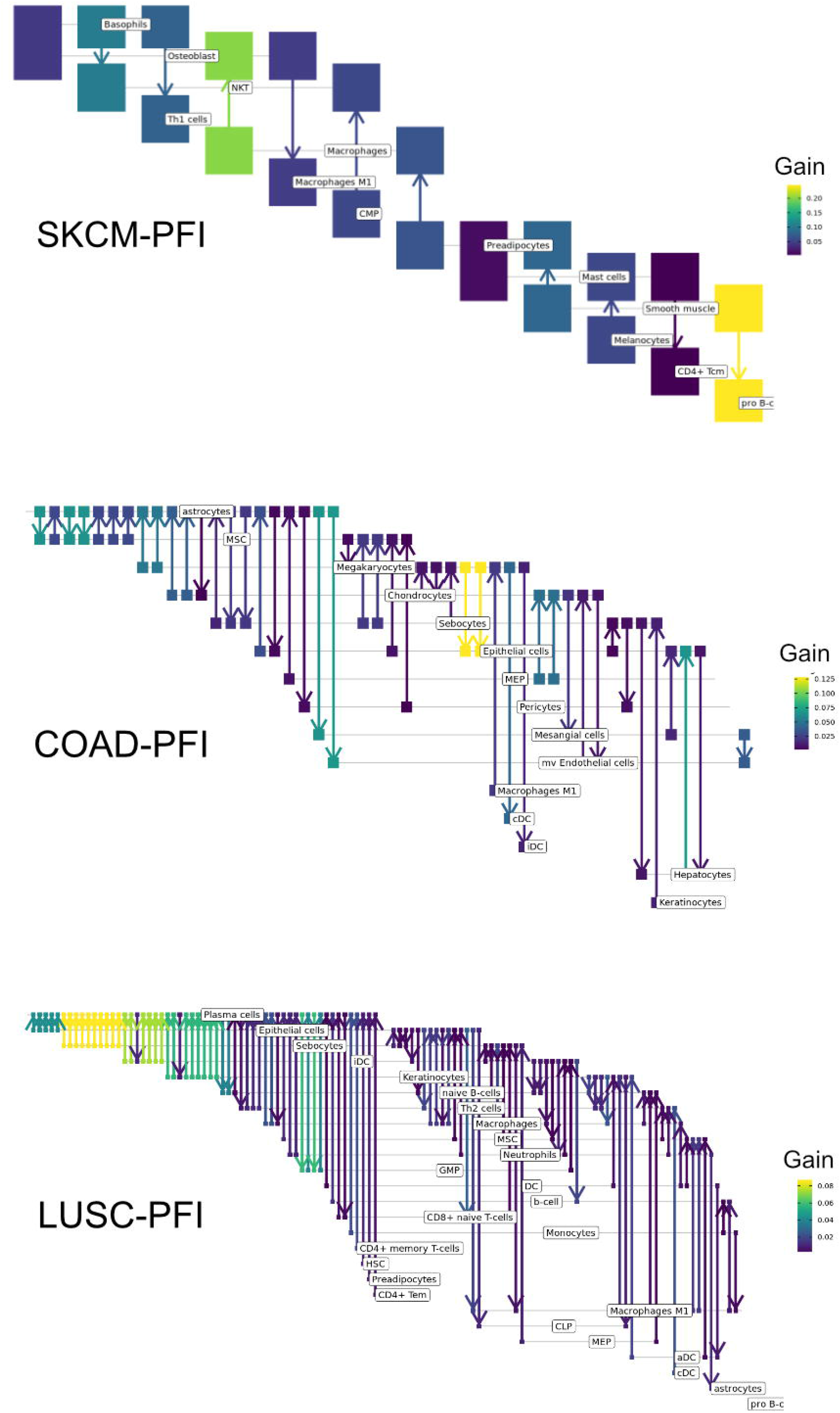
Informative edges selected by XGBoost models for prediction within study. Color indicates information gain.

The purpose of the modeling was within-data inference rather than classification outside of the TCGA pan-cancer data set. After fitting, it is possible to examine what model features (edges) are most useful in classification. The XGBoost classifiers are regularized models, not all features will be used and often only a small subset of features are retained in the final model. We assess the relative usefulness of a feature by comparing the feature gain -- the improvement in accuracy when a feature is added to a tree.

All classification models had an accuracy between 91% (SKCM, PFI) and 99% (COAD, Stage). As mentioned above, there can be a high degree of correlation between edge values in a data set. While features are selected first based on improving prediction, the machine learning model accounts for correlated features by selecting the one that has best predictive power, leaving out other correlated features. That said, the number of features selected by the model is then related to the correlation structure. In a set of uncorrelated features where all features add to the predictive power, all features will be selected, whereas for correlated features, only a small number will be selected. This is seen in results here in terms of differences in the numbers of features compared with the starting network.

In the COAD-PFI case, the number of features was reduced by approximately 20%, keeping 50 edges in the model. The STAD-PFI features were reduced by approximately 45%. Other examples are are LUSC-PFI at 60% reduction, ESCA-PFI at 74%, and SKCM-PFI at 95% (12 edges selected) indicating a high degree of internal feature correlation.

A similar pattern was observed in the tumor stage contrasts, where SKCM-stage had a 96% reduction in features, STAD-stage 52%, READ-stage 47%. For COAD-stage, feature reduction was 95% reduction, but attributable to the large number of starting edges (1851) compared to the 84 edges selected. A collection of the most predictive edges is given in Table 2.

**Table 2.** Top 5 most predictive edges from XGBoost models.

| Contrast | Study | EdgeID | LCell | Ligand | Receptor | RCell | S1 | Median Diff | Information Gain |
| --- | --- | --- | --- | --- | --- | --- | --- | --- | --- |
| PFI | COAD | 586640 | Megakaryocytes | BMP10 | ENG | Epithelial cells | 0.169 | 0.082 | 0.109 |
| PFI | COAD | 50871 | astrocytes | TNC | ITGA5 | mv Endothelial cells | 0.168 | 0.061 | 0.069 |
| PFI | COAD | 406871 | Hepatocytes | GDF2 | ENG | Epithelial cells | 0.168 | 0.082 | 0.067 |
| PFI | COAD | 49669 | astrocytes | EFNB1 | EPHB4 | Mesangial cells | 0.199 | 0.117 | 0.066 |
| PFI | COAD | 632560 | MEP | TIMP2 | ITGB1 | MEP | 0.167 | 0.095 | 0.051 |
| Stage | COAD | 406579 | Hepatocytes | CGN | TGFBR2 | Eosinophils | -0.165 | -0.077 | 0.043 |
| Stage | COAD | 330377 | Eosinophils | LAMB3 | ITGB1 | Eosinophils | -0.144 | -0.048 | 0.038 |
| Stage | COAD | 616033 | Memory B-cells | BMP15 | BMPR2 | Epithelial cells | -0.150 | -0.060 | 0.037 |
| Stage | COAD | 784400 | NK cells | TNFSF10 | TNFRSF10B | CD4+ memory T-cells | 0.137 | 0.043 | 0.037 |
| Stage | COAD | 630108 | MEP | B2M | KIR2DL1 | iDC | 0.138 | 0.055 | 0.037 |
| PFI | ESCA | 457801 | Keratinocytes | GS | ADCY7 | CD4+ Tcm | 0.167 | 0.078 | 0.078 |
| PFI | ESCA | 182483 | CD8+ Tcm | RBP3 | NOTCH1 | pDC | -0.184 | -0.073 | 0.071 |
| PFI | ESCA | 1051114 | Th2 cells | CALM1 | GP6 | naive B-cells | 0.171 | 0.085 | 0.070 |
| PFI | ESCA | 658080 | Mesangial cells | SPP1 | CD44 | Tregs | 0.184 | 0.080 | 0.064 |
| PFI | ESCA | 397215 | GMP | HMGB1 | THBD | MEP | 0.184 | 0.060 | 0.059 |
| PFI | LUSC | 879775 | Plasma cells | VEGFA | ITGB1 | GMP | 0.120 | 0.047 | 0.041 |
| PFI | LUSC | 451902 | iDC | VEGFA | ITGB1 | Plasma cells | 0.137 | 0.067 | 0.038 |
| PFI | LUSC | 398971 | GMP | ADAM17 | ITGB1 | Plasma cells | 0.120 | 0.059 | 0.030 |
| PFI | LUSC | 340857 | Epithelial cells | COL4A6 | ITGB1 | CD8+ naive T-cells | 0.124 | 0.054 | 0.026 |
| PFI | LUSC | 471558 | Keratinocytes | THBS1 | ITGA6 | Plasma cells | 0.120 | 0.068 | 0.025 |
| Stage | READ | 632552 | MEP | TGFB3 | TGFBR2 | MEP | -0.267 | -0.134 | 0.127 |
| Stage | READ | 795527 | NKT | GZMB | PGRMC1 | CD4+ memory T-cells | 0.343 | 0.144 | 0.115 |
| Stage | READ | 402754 | Hepatocytes | CGN | TGFBR2 | CD4+ Tem | -0.274 | -0.134 | 0.108 |
| Stage | READ | 808308 | NKT | GZMB | IGF2R | Plasma cells | 0.261 | 0.101 | 0.103 |
| Stage | READ | 800747 | NKT | IL7 | IL2RG | GMP | 0.264 | 0.136 | 0.095 |
| PFI | SKCM | 1008243 | Smooth muscle | SEMA7A | PLXNC1 | pro B-cells | 0.438 | 0.259 | 0.242 |
| PFI | SKCM | 517677 | Macrophages | UBA52 | NOTCH1 | Osteoblast | -0.284 | -0.145 | 0.200 |
| PFI | SKCM | 80934 | Basophils | VIM | CD44 | NKT | -0.383 | -0.254 | 0.103 |
| PFI | SKCM | 1007915 | Smooth muscle | PSAP | SORT1 | Preadipocytes | 0.311 | 0.175 | 0.082 |
| PFI | SKCM | 84049 | Basophils | CALM1 | PTPRA | Th1 cells | -0.285 | -0.151 | 0.080 |
| Stage | SKCM | 275306 | CLP | GI2 | CXCR1 | Osteoblast | 0.353 | 0.176 | 0.207 |
| Stage | SKCM | 399084 | GMP | TIMP1 | CD63 | Plasma cells | -0.302 | -0.147 | 0.206 |
| Stage | SKCM | 273727 | CLP | GI2 | F2R | MEP | 0.290 | 0.123 | 0.182 |
| Stage | SKCM | 182981 | CD8+ Tcm | GI2 | TBXA2R | Plasma cells | -0.283 | -0.095 | 0.123 |
| Stage | SKCM | 397545 | GMP | BST1 | CAV1 | MSC | -0.337 | -0.194 | 0.109 |
| PFI | STAD | 461765 | Keratinocytes | CALM3 | KCNQ1 | Eosinophils | -0.136 | -0.067 | 0.062 |
| PFI | STAD | 644724 | Mesangial cells | TGFB2 | ACVR1 | Erythrocytes | 0.149 | 0.061 | 0.054 |
| PFI | STAD | 105991 | CD4+ T-cells | IL1B | IL1R2 | Megakaryocytes | 0.134 | 0.081 | 0.047 |
| PFI | STAD | 269013 | CLP | ADAM28 | ITGA4 | CD4+ T-cells | 0.145 | 0.075 | 0.046 |
| PFI | STAD | 343620 | Epithelial cells | VCAN | TLR1 | CLP | 0.134 | 0.051 | 0.033 |
| Stage | STAD | 128412 | CD4+ Tem | CALM1 | KCNQ1 | Macrophages | 0.140 | 0.058 | 0.057 |
| Stage | STAD | 43832 | astrocytes | FBN1 | ITGB6 | Epithelial cells | -0.146 | -0.058 | 0.036 |
| Stage | STAD | 346120 | Epithelial cells | LAMB1 | ITGAV | Hepatocytes | -0.139 | -0.066 | 0.035 |
| Stage | STAD | 403540 | Hepatocytes | SHH | PTCH1 | CD8+ T-cells | -0.138 | -0.069 | 0.034 |
| Stage | STAD | 648983 | Mesangial cells | FGB | ITGAV | Megakaryocytes | -0.140 | -0.060 | 0.031 |

Contrast: the groupwise test performed, Study: tissue type, Edge ID: BigQuery table lookup ID, LCell: cell producing ligands, Ligand: ligand gene symbol, Receptor: receptor gene symbol, R Cell: receptor producing cell, $S_1$ : between group $S_1$ statistic, Median Diff: difference in edge weights between groups, Information Gain: Xgboost information gain after adding feature to model.

The collection of genes from each differential network was used for gene ontology (GO) term enrichment using the GONet tool (Pomaznoy et al., 2018). All tissue-contrast combinations with differentially weighted edges produced enriched GO terms (FDR < 0.05, within tissue contrasts) except the SKCM-stage group, which although contained 77 genes in the differential network, produced no enriched terms.

Common themes included structural GO terms such as “extracellular structure organization” (for SKCM), cell-substrate adhesion (ESCA, LUSC), cell-cell adhesion (STAD), ECM / extracellular matrix organization (LUSC, COAD, READ, STAD). Cell migration was also a common theme with “cell migration” (STAD), “epithelial cell migration” (SKCM), and “regulation of cell migration” (LUSC, COAD/READ). Among immune related themes, GO terms included “IFNG signaling” and “antigen processing and presentation” (SKCM), “regulation of immune processes” and “IL2” (STAD), and “viral host response” (COAD / READ). See Table 3 for a summary and supplemental table 3 for complete results.

**Table 3.** Enriched GO terms.

| <u>Tissue</u> | <u>Contrast</u> | <u>Num GOs</u> | <u>ECM</u> | <u>Migration</u> | <u>Immune</u> | <u>Immune2</u> |
| --- | --- | --- | --- | --- | --- | --- |
| SKCM | PFI | 34 | extracellular structure organization | epithelium cell migration | IFNG signaling | antigen processing and presentation |
| ESCA | PFI | 3 | cell-substrate adhesion |  |  |  |
| STAD | PFI | 59 | cell-cell<br>adhesion<br>mediated by<br>integrin | cell migration | regulation of<br>immune<br>system process | IL2 |
| LUSC | PFI | 39 | extracellular<br>matrix<br>organization | positive<br>regulation of<br>cell migration |  |  |
| COAD / READ | stage | 85 | ECM | regulation of<br>epithelial cell<br>migration | viral host<br>response |  |
| STAD | stage | 28 | ECM /<br>adhesion | cell migration |  |  |

Tissue: TCGA study, Contrast: the groupwise test performed, Num GOs: number of gene ontology terms found significantly enriched, ECM: GO categories involving ECM, Migration: GO terms involving cell migration, Immune: GO terms involving immune response, Immune2: additional GO terms involving immune response.

## Discussion

Patient outcome or response to therapy is not necessarily well predicted by tumor stage alone (Kirilovsky et al., 2016). As Fridman et al. wrote, “different types of infiltrating immune cells have different effects on tumour progression, which can vary according to cancer type” (Fridman et al., 2012). This idea has been developed further with the creation of the ‘Immunoscore’, a prognostic based on the presence and density of particular immune cells in the TME context, expanded to include the peripheral margin as well as tumor core. For example, the Immunoscore in colorectal cancer depends on the density of both CD3+ lymphocytes (any T cell) and specifically, CD8+ cytotoxic T cells in the tumor core and invasive margin (Pagès et al., 2018). The differences in factors that relate to stage and survival is reflected in the current work in the identification of different cell-cell interactions of importance for each.

Previous studies have shown that cellular interactions within the tumor microenvironment have an impact on patient survival, drug response, and tumor growth. X. Zhao et al. (Zhou et al., 2017) described alterations in ligand-receptor pair associations in cancer compared to normal tissue, the cell-cell communication structures thereby becoming a generalized phenotype for malignancy. Using the same foundational database of possible interactions as this work, ligand-receptor pair expression correlation was compared between tumor and normal tissue. Their “aggregate analysis revealed that … tumors of most cancer types generally had reduced (ligand-receptor) correlation compared with the normal tissues.” The ligand-receptor pairs that commonly showed such differences across the ten tissue types studied included PLAU-ITGA5, LIPH-LPAR2, SEM14G-PLXNB2, SEMABD-TYROBP, CCL2-CCR5, CCL3-CCR5, and CGN-TYROBP.

Like the Zhao et al. work, we found the collection of associated edges enriched for related biological processes, especially to ECM organization and cell adhesion -- possibly related to the progression towards dysplasia. For example, in Zhao et al., the ligand-receptor pairs COL11A1-ITGA2, COL7A1-ITGA2, MDK-GPC2 and MMP1-ITGA2 were found to be positively correlated in cancer but not in normal tissue. In the current work, integrins and laminins generally have elevated edge weights in late tumor stage. In the PFI contrasts, except for ESCA, such edges have higher weights in shorter PFIs, corresponding to more severe progression. Regarding SEMA7A, found in the PFI STAD results as a predictive feature, previous findings report the collagen gene COL1A1 has been associated with metastasis, and SEMA7A is known to play an important role in integrin-mediated signaling and functions both in regulating cell migration and immune responses. Cancers such as esophageal, gastric, and colorectal all show transitions to metaplasia and dysplasia, a process that breaks down the structural order of a tissue, replacing it with disorder and cell transdifferentiation.

In our model, a host response is reflected in a change in *S_1_* score, negative if the edge weight is higher with longer PFI times. In the PFI results, Th1 cells appeared in 13 high scoring edges in SKCM, all with negative *S_1_* values. Also, for SKCM and COAD, ligand producing (pro-inflammatory) M1 macrophage edges are present but show both positive and negative *S_1_* scores. Inflammation cytokines IL1B and IL18 are both present in the results of ESCA and STAD (Figure 9). In the tumor stage contrasts, we see Th2 and NK cells with inflammation cytokines IL1A, IL1B, IL4, TNF in STAD and COAD. So, while certain inflammatory signatures are observed, the absence of well-known canonical edges such as Th1-IL12-IL12RB1-M1 macrophages, may be due to essentially no difference, or undetectable differences in the quantity of Th1 cells or IL12A expression between PFI groups (4.9 vs 3.3 TCGA Pan-Cancer RSEM for short vs long PFI). These observations point to possible mechanisms of action for immune cells known to be important for cancer immune response, the CD4+ T helper 1 cells and M1 macrophages, in relation to tumor progression

**Figure 9.**
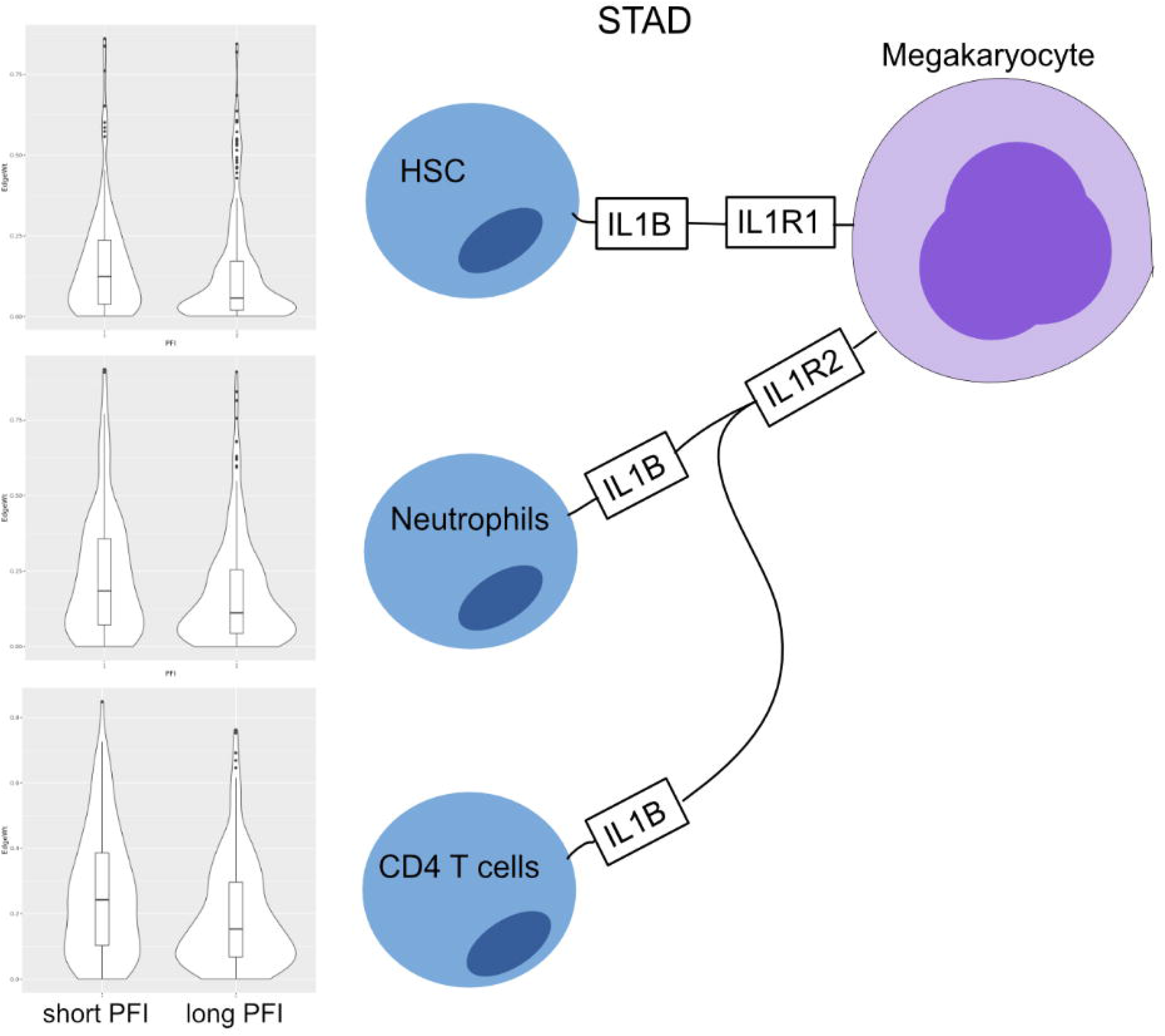
Cell-cell interaction diagram demonstrating complexity in communication with three cell types that produce the IL1B ligand that have two possible binding partners on the same receptor bearing cell. Edge weight violin plots are shown for two STAD PFI groups, short (left) and long (right) PFI.

In tissues susceptible to dysplasia, such as the tissues explored here, unexpected cell types may be detected. For example, the ‘…disruption of tissue organization appears to trigger a profound change in cellular commitment, which leads to hepatocyte differentiation in the “oval cells” in … the epithelial cells lining the small pancreatic ductules’ (Reddy et al., 1991). As another example, pancreatic cancer is known to have desmoplastic stroma, the source of which may include MSCs which are defined by their ability to differentiate into osteoblasts, chondrocytes, and adipocytes (Mathew et al., 2016). In line with that finding, it’s been observed that “…stromal cells isolated from the neoplastic pancreas can differentiate into osteoblasts, chondrocytes, and adipocytes” (Mathew et al., 2016).

It has been reported that (Yáñez et al., 2017), “granulocyte-monocyte progenitors (GMPs) and monocyte-dendritic cell progenitors (MDPs) produce monocytes during homeostasis and in response to increased demand during infection.” or as in (Weston et al., 2018), “Granulocyte-monocyte progenitor (GMP) cells play a vital role in the immune system by maturing into a variety of white blood cells, including neutrophils and macrophages, depending on exposure to cytokines such as various types of colony stimulating factors (CSF).”

In our results for SKCM and COAD, GMPs had negative S1 statistics, meaning the late-stage cases had edges with higher weights. The GMP cells most often interacted with (as receptor bearing cells) MSC, Melanocytes, both M1 and M2 macrophages, and CD8+ Tem (T effector-memory cells). The presence of GMP related edges may be indicative of the commonly observed ‘myeloid dysfunction’, which “can promote tumor progression through immune suppression, tissue remodeling, angiogenesis or combinations of these mechanisms.“(Messmer et al., 2015) Also, “tumors secrete a variety of factors such as G-CSF that act in a systemic way to reduce IRF-8 within progenitor cells, releasing myelopoiesis from IRF-8 control such that the granulocytic lineage (blue cell) undergoes hyperplasia, leading to increased immature suppressive cells to promote tumor growth.” This is in line with our observations.

Megakaryocytes, a multipotent stem cell, are cells that typically reside in the bone marrow and produce platelets. Megakaryocytes are also produced in the liver, kidney, and spleen. Additionally, megakaryocytes have been observed in the lung and circulating blood where they were useful as a biomarker in prostate cancer. Case reports exist showing megakaryocytes in the metaplasia of gastric cancer patients (Chatelain et al., 2004). Megakaryocytes respond to a variety of cytokines such as IL-3, IL-6, IL-11, CXCL5, CXCL7, and CCL5. A majority of interacting cells are leukocytes. In both esophageal and gastric cancers “…thrombocytosis has been reported in general to be associated with adverse clinical outcomes. (Voutsadakis, 2014)” Additionally, there are reports of ‘tumor educated platelets’ that can be useful as part of a liquid biopsy (Best et al., 2018) (Haemmerle et al., 2018).

Among the rich literature regarding oncological cytokine networks, there is a strong emphasis on the cancer cell as a central actor. Many of the review articles and research focuses on the cancer cell interactions in the TME. For example, cancer cells producing an overabundance of IL6 or IL10 that has been associated with poor prognosis (Burkholder et al., 2014; Fisher et al., 2014; Lippitz and Harris, 2016).

However, in this work, the focus has been put on the environment and less about the cancer cell itself. This is largely because in performing cell deconvolution on gene expression data to determine the presence and quantity of different cell types in the mixed sample, reliable signatures for cancer cells are not readily available. Because in carcinomas, a cancer cell derives from the epithelium, and in many ways remains similar to epithelial cells. Even in single cell RNA-seq studies, it is often difficult to determine what cells are cancerous and picking this signature out of a mixed expression dataset is difficult and remains an open question.

This work is based upon gene expression, rather than protein expression, cell-surface expression or secretion measurements. Also, the base expression data is taken from sorted cells, rather than cells in tissue with an assumption that we cannot get “new/non-scaffold edges” in a tissue/cancer context.

However new data types and methods including scRNA-seq and PIC-seq will provide ways of determining new cell-cell interactions that are context specific (Giladi et al., 2020).

Importantly, the physical and biochemical process of secretion, binding and activation cannot be identified with the current data and method. By identifying the propensity of edge constituents in particular tumor microenvironments in comparison with others, it becomes more likely that communication with activation can take place, as the presence of those constituents is a prerequisite.

With the data and results publicly available in a Google BigQuery table (Supplemental Figure 8), this resource is open to researchers to explore and ask questions. It is a low-cost way (with free options) to achieve compute cluster performance for quickly answering such questions. The table is easily joined to clinical and molecular annotations and can be worked with from R and python notebooks. With the addition of resources like GTEx, it should begin to be possible to tease aberrant, cancer specific interactions apart.

In terms of future work, it could be important to examine communication networks given the immune subtypes of (Thorsson et al., 2018) and communication differences between TCGA tumor molecular subtypes. New data types can be applied to enhance the scaffold with knowledge gained from (for example) single-cell RNA-seq.

In this work, we have introduced a method and identified lines of communication between cells that may play a role in disease. These lines include both established/recognized cells in the context of cancer, as well as others that should be explored further, with targeted methods.

## Acknowledgments

The authors would like to thank Samuel Danziger, David Reiss, Mark McConnell, Andrew Dervan, Matthew Trotter, Douglas Bassett, Robert Hershberg, the Shmulevich Lab and the Institute for Systems Biology for engaging and informative discussions. This study was supported by Celgene, a wholly owned subsidiary of Bristol-Myers Squibb, in part through a Sponsored Research Award to D.L.G., B.A. and I.S, and by the Cancer Research Institute (D.L.G, V.T, I.S). We thank the ISB-CGC for their ongoing support. ISB-CGC has been funded in whole or in part with Federal funds from the National Cancer Institute, National Institutes of Health, Task Order No. 17X148under Contract No. 75N91019D00024. The content of this publication does not necessarily reflect the views or policies of the Department of Health and Human Services, nor does mention of trade names, commercial products, or organizations imply endorsement by the U.S. Government.

## Author Contributions

D.L.G., B.A., V.T, A.R., I.S. conceived of the idea. D.L.G. developed the method, wrote the code, and performed the computations. D.L.G. wrote the manuscript with contributions from B.A., V.T., A.R., I.S. and A.R. supervised the project. All authors provided critical feedback and helped shape the research, analysis and manuscript.

