## Supplemental Figures for "Patient-specific cell communication networks associate with disease progression in cancer"

### Abstract

The maintenance and function of tissues in health and disease depends on cell-cell communication. This work shows how high-level features, representing cell-cell communication, can be defined and used to associate certain signaling 'axes' with clinical outcomes. Using cell-sorted gene expression data, we generated a scaffold of cell-cell interactions and define a probabilistic method for creating per-patient weighted graphs based on gene expression and cell deconvolution results. With this method, we generated over 9,000 graphs for TCGA patient samples, each representing likely channels of intercellular communication in the tumor microenvironment . It was shown that particular edges were strongly associated with disease severity and progression, in terms of survival time and tumor stage. Within individual tumor types, there are predominant cell types and the collection of associated edges were found to be predictive of clinical phenotypes. Additionally, genes associated with differentially weighted edges were enriched in Gene Ontology terms associated with tissue structure and immune response. Code, data, and notebooks are provided to enable the application of this method to any expression dataset (https://github.com/IlyaLab/Pan-Cancer-Cell-Cell-Comm-Net).

Supplemental Figure 1

Edge weight distribution

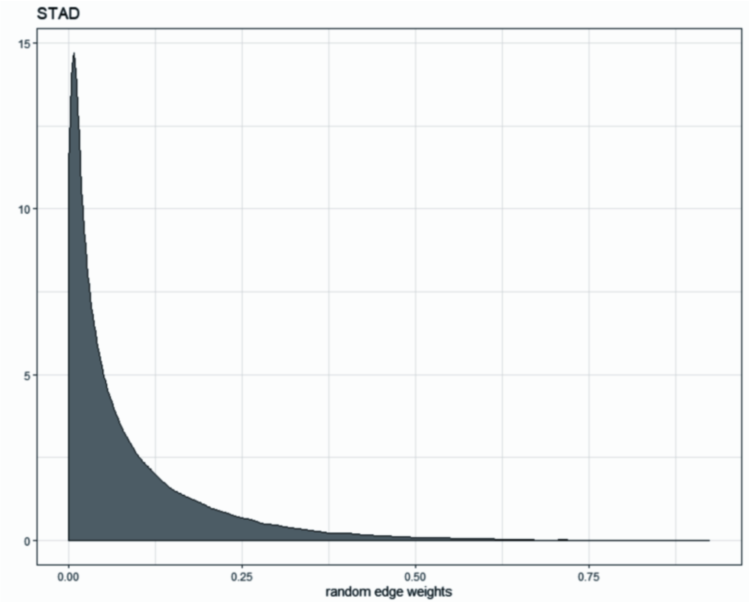


Distribution of edge weights for STAD samples over a random sample of edges.

Resampled $S_1$ statistic distributions

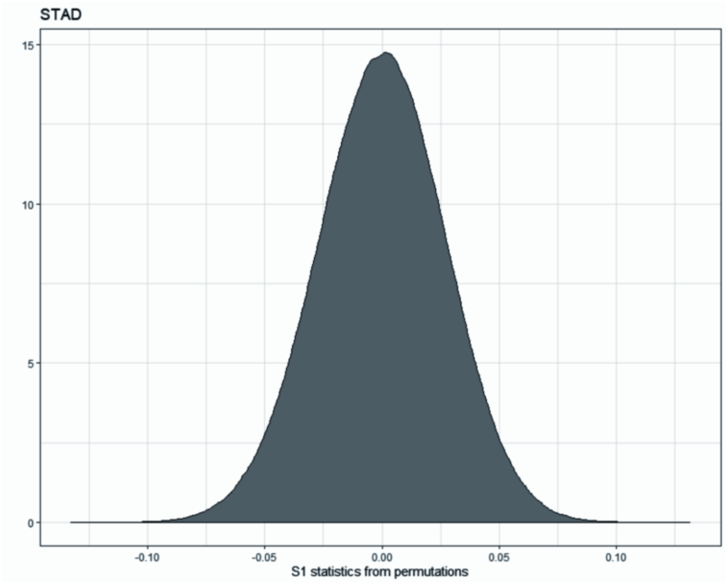


Example of distribution of $S_1$ statistics for STAD samples after permuting edge weights.

### Supplemental Figure 2

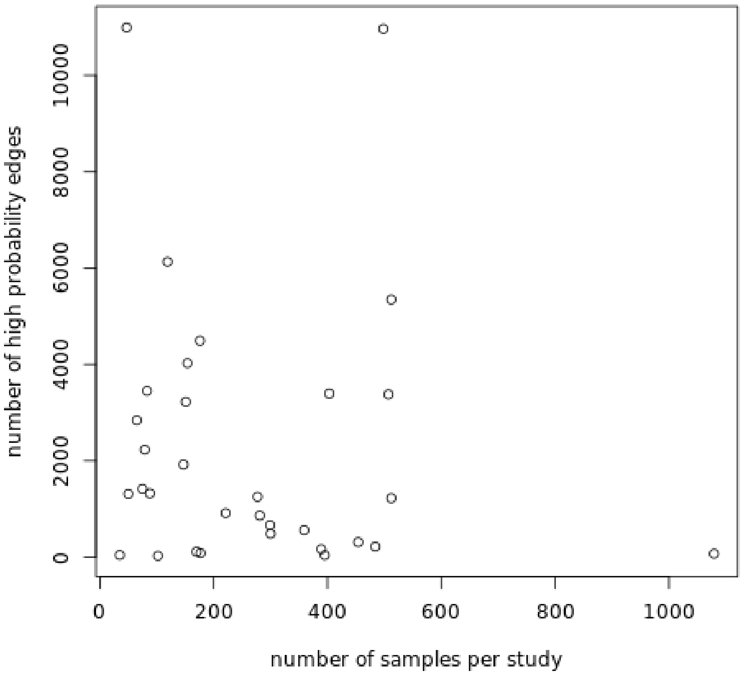


Study tests show number of high probability edges not strongly associated with number of samples.

Supplemental Figure 3

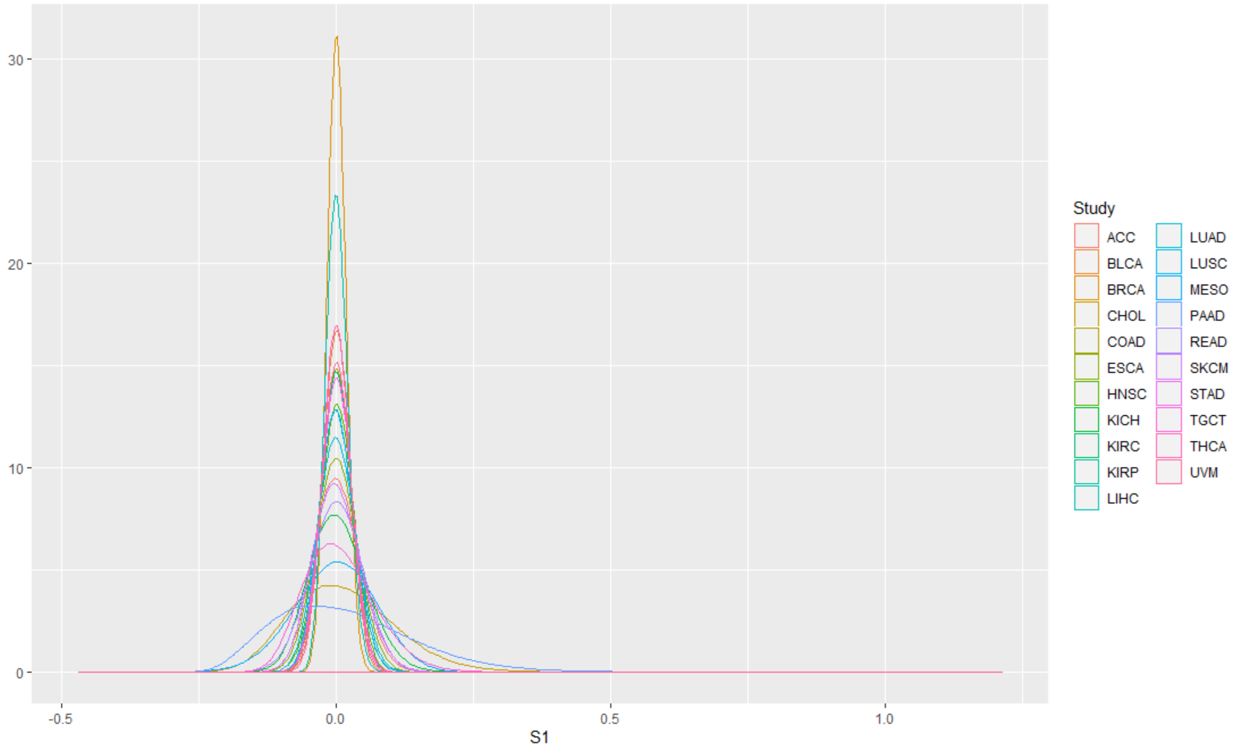


Resampled $S_1$ statistics distributions within study for early vs late stage phenotypes. Permuted sample sizes matched sample sizes for actual tests.

Supplemental Figure 4

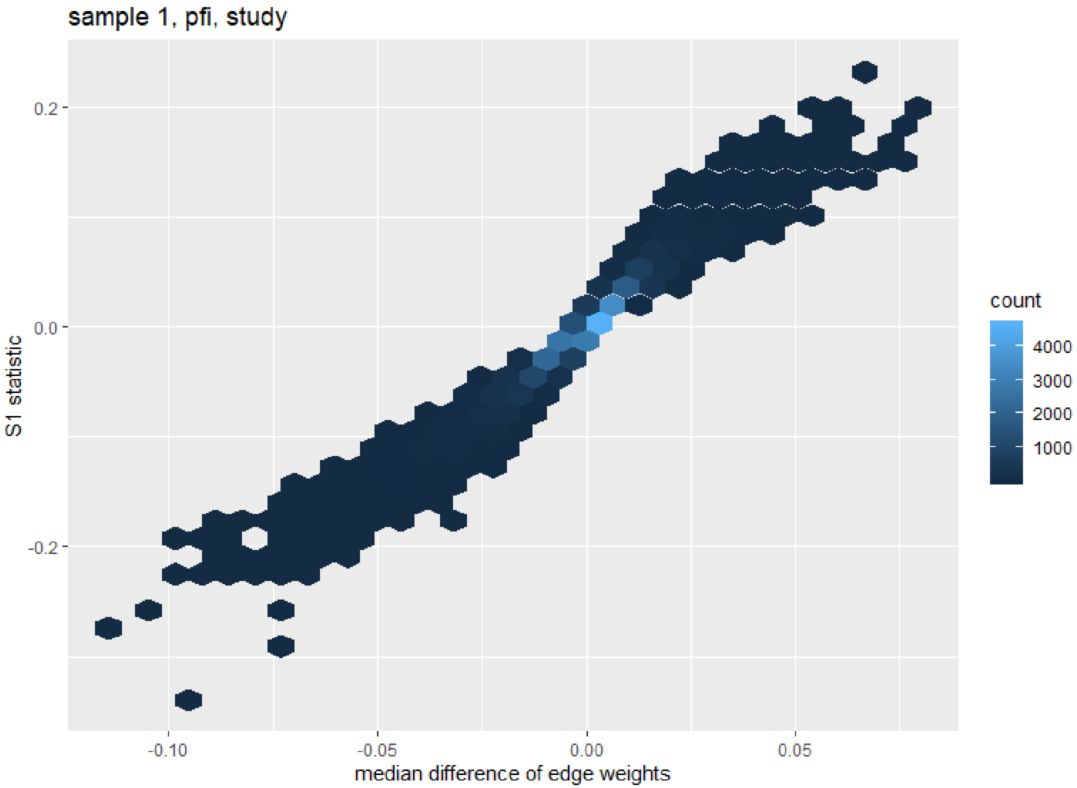


S1 statistics scale with median differences of edge weights.

### Supplemental Figure 5

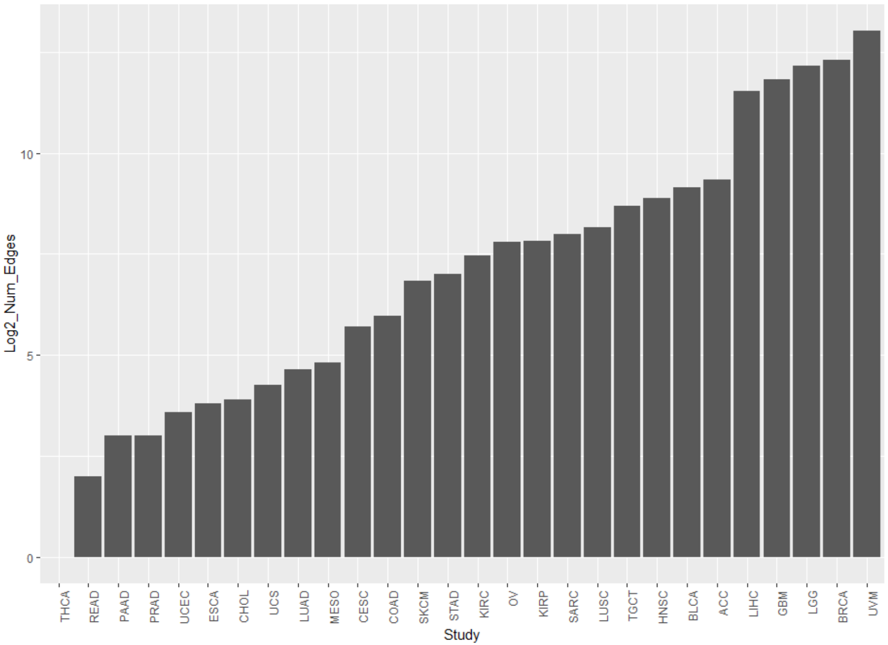


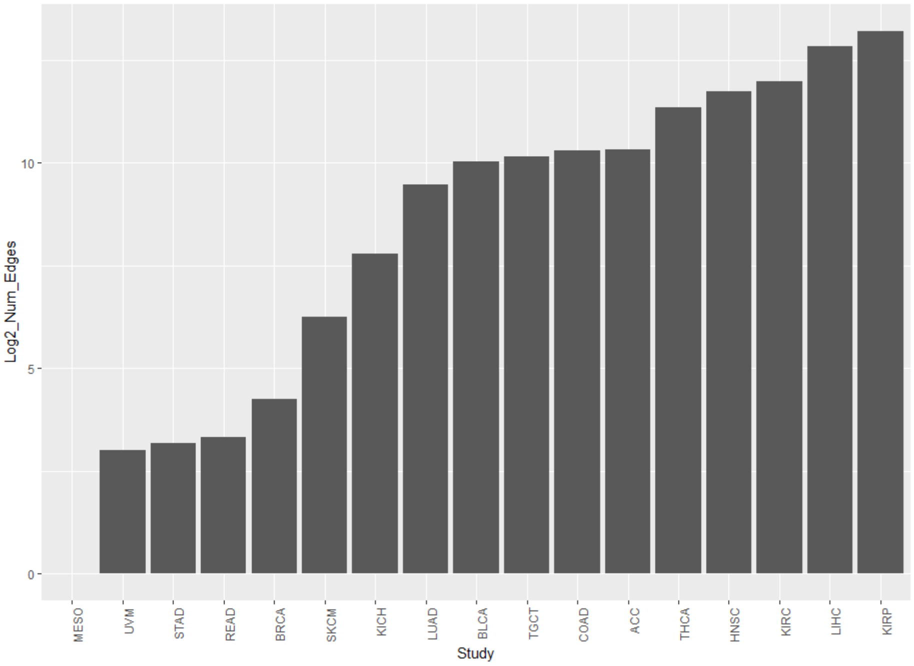


Study - PFI results, counts of edges that have $S_1$ statistics beyond the 1 millionth% of permuted results.

Study - Stage results, counts of edges that have $S_1$ statistics beyond the 1 millionth percent of permuted results.

Supplemental Figure 6A

Study - PFI, STAD

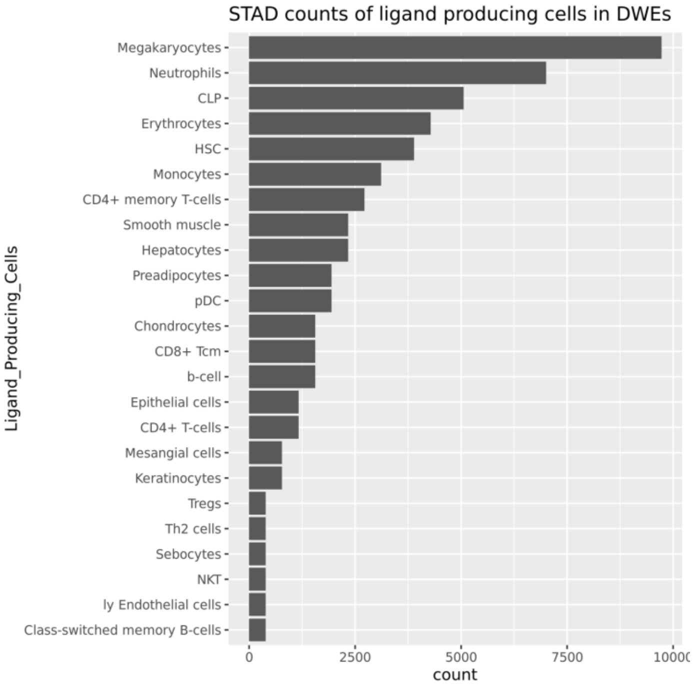


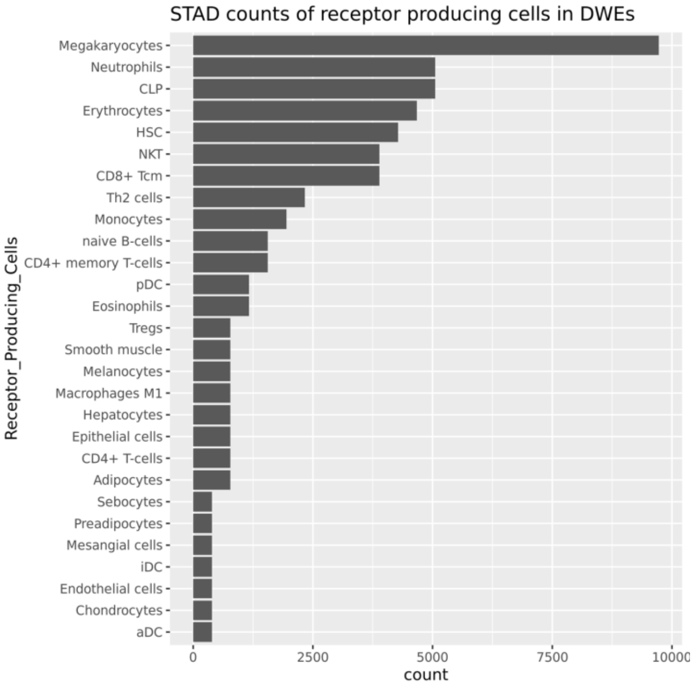


Edges with high statistics under a given phenotype contrast are dominated by specific cell types.

Supplemental Figure 6B

Study-PFI, STAD, Ligands

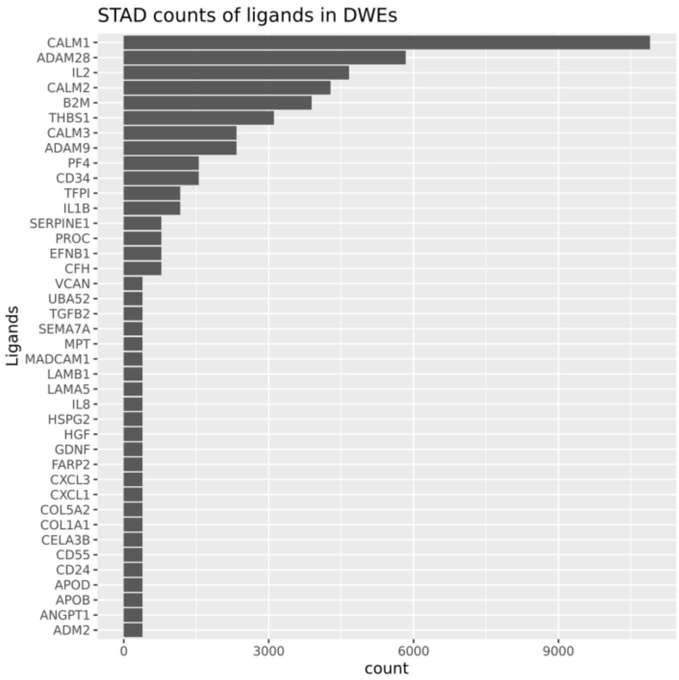


Study-PFI, STAD, Receptors

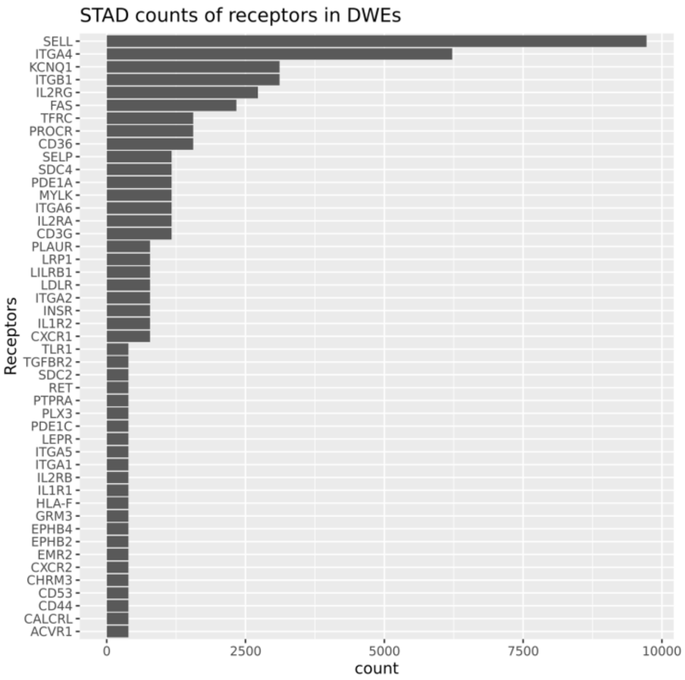


Common components (top 50) in high statistic edges for the STAD-PFI contrast.

### Supplemental Figure 7

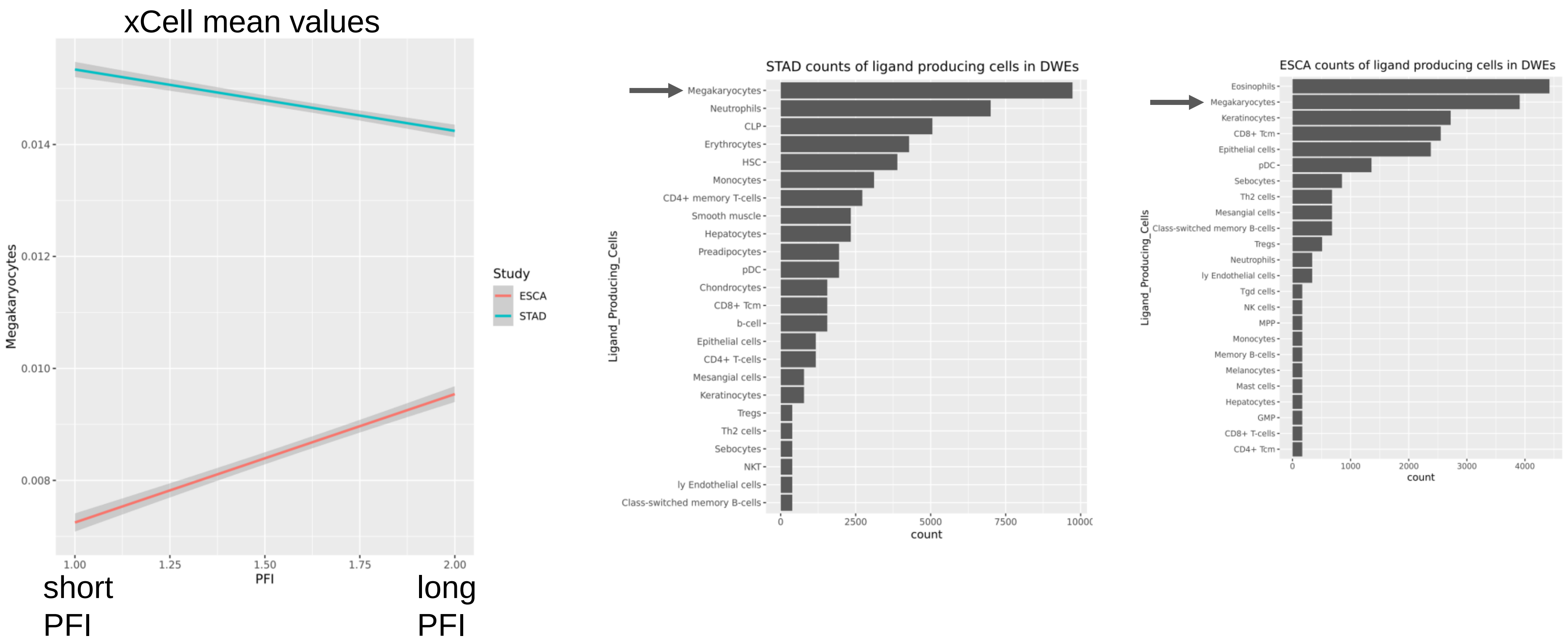


Megakaryocytes decrease with PFI in STAD, but increase in ESCA, and are part of many of the differentially weighted edges (DWEs).

#### Supplemental Figure 8

Google BigQuery database sanity check. Queried for edge weights for STAD, in the high probability result edges (in github also). Top hit was edge-ID 597043 (as indexed in BigQuery). Results listed median difference as 0.14. Manual computation gives the same result.

| | | | | | | $S_1$ | Med. Diff. |
|---|---|---|---|---|---|---|---|
| STAD | 597043 | Megakaryocytes | IL2 | IL2RG | Th2 cells | 0.24188105422239894 | 0.1387622086 |

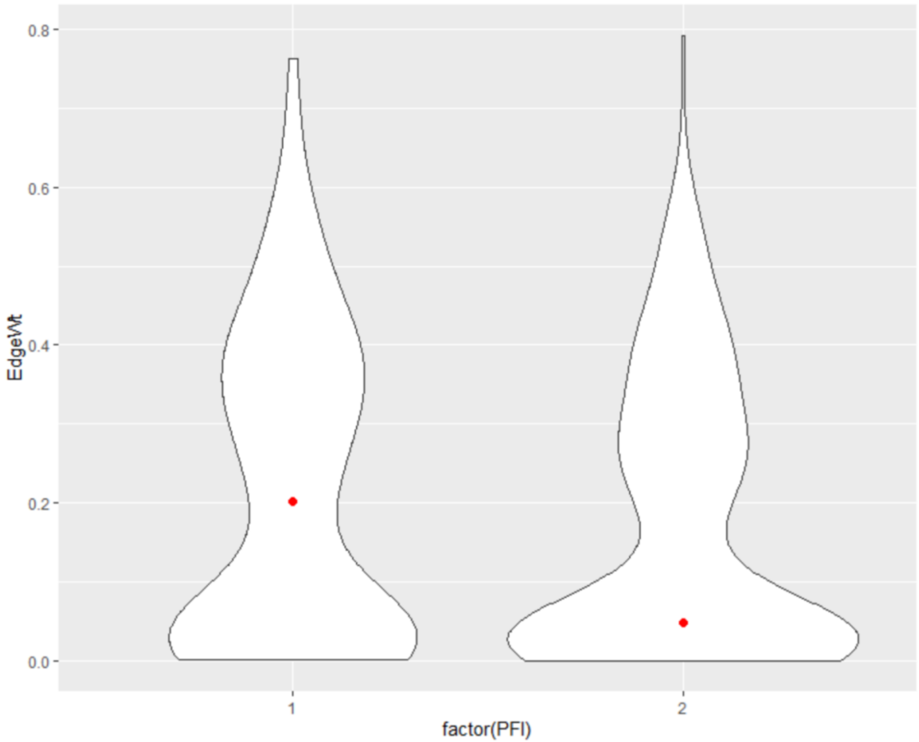


Red dots are the medians given these edge weights.
